# Regenerating insulin-producing β-cells ectopically from a mesodermal origin in the absence of endothelial specification

**DOI:** 10.1101/2021.02.01.429189

**Authors:** Ka-Cheuk Liu, Alethia Villasenor, Nicole Schmitner, Niki Radros, Linn Rautio, Sven Reischauer, Didier Y.R. Stainier, Olov Andersson

## Abstract

To investigate the role of the vasculature in pancreatic β-cell regeneration, we crossed a zebrafish β-cell ablation model into the avascular *npas4l* mutant (i.e. *cloche*). Surprisingly, β-cell regeneration increased markedly in *npas4l* mutants owing to the ectopic differentiation of β-cells in the mesenchyme, a phenotype not previously reported in any models. The ectopic β-cells expressed endocrine markers of pancreatic β-cells, and also reduced glucose levels in the β-cell ablation model. Through lineage tracing, we determined that the vast majority of these ectopic β-cells derived from the mesodermal lineage. Notably, ectopic β-cells were found in *npas4l* mutants as well as following knockdown of the endothelial determinant Etv2. Together, these data indicate that in the absence of endothelial specification, mesodermal cells possess a remarkable plasticity enabling them to form β-cells, which are normally endodermal in origin. Understanding the restriction of this differentiation plasticity will help exploit an alternative source for β-cell regeneration.

## Introduction

The concept of embryonic development and cell fate determination was illustrated by the famous Waddington landscape model decades ago (Waddington, 1957). Waddington’s model not only shows the importance of spatiotemporal precision in cell differentiation but also metaphorizes cell fate determination as a sequential and irreversible event. In this hierarchical model, endoderm follows the lineage paths downwards and progressively differentiates into multiple endodermal cell types, including pancreatic β-cells. Likewise, mesoderm stays in the mesodermal lineage paths and differentiates into vasculature and other mesodermal cell types. However, in recent decades, multiple studies have suggested that committed cells are capable of differentiating across the germ layer border by converting embryonic and/or adult mesodermal fibroblasts into ectodermal neuronal cells (Vierbuchen et al., 2010), multipotent induced neural stem cells (Ring et al., 2012), endodermal hepatocyte-like cells (Huang et al., 2011; Sekiya & Suzuki, 2011) or pancreatic β-like cells (Zhu et al., 2016) *in vitro*. These studies highlight the feasibility of converting mesodermal cells into ectodermal or endodermal cells *in vitro* after the addition of factors.

Despite the extensive studies on cell fate conversion across germ layers *in vitro*, the number of *in vivo* studies is limited. Ectopic expression of *Xsox17β* in *Xenopus* embryos relocated cells normally fated for ectoderm to appear in the endodermal gut, suggesting a possible change in cell fate *in vivo* (Clements & Woodland, 2000). Furthermore, aggregated morulae and chimeric embryos of β-catenin mutants provided evidence of precardiac mesoderm formation in the endodermal region *in vivo* (Lickert et al., 2002). Unlike studies expressing ectopic transcription factors or inducing mutations, the study by Goldman and collaborators revealed endodermal cells differentiating into endothelial cells, which were believed to be mesodermal derivatives, during normal liver development in lineage-tracing mouse models (Goldman et al., 2014). These studies suggest that the classical *in vivo* germ layer border may not be as clear-cut as previously thought.

In this study, we aimed to elucidate the importance of the vasculature in pancreatic β-cell regeneration, which plays a crucial role in potential therapeutic strategies against diabetes. We employed *cloche* zebrafish mutants as an avascular model. The mutation of *npas4l*, a master regulator of endothelial and hematopoietic cell fates, is responsible for the severe loss of most blood vessels and blood cells in *cloche* mutants (Parker & Stainier, 1999; Reischauer et al., 2016; Stainier et al., 1995). Unexpectedly, the *npas4l* mutation induced ectopic β-cell formation in the mesenchymal region outside the pancreas and decreased the glucose level after β-cell ablation. Lineage-tracing mesodermal cells expressing *draculin* (*drl*) and *etv2* validated the mesodermal lineage of the ectopic β-cells, which are normally endodermal in origin. These findings offer novel insights into cell fate determination and an alternative source of β-cells.

## Results

### Ectopic β-cell formation and improved glucose control in *npas4l* mutants

To determine the importance of vasculogenesis and vascularization for β-cell regeneration, we examined β-cell formation in zebrafish carrying the *cloche* mutation (*npas4l*^−/−^) after β-cell ablation, i.e., in the *Tg(ins:Flag-NTR);Tg(ins:H2BGFP;ins:DsRed)* model. Nitroreductase (NTR), expressed by the *insulin* promoter, converts the prodrug metronidazole (MTZ) to a cytotoxin to specifically ablate insulin-producing β-cells (Curado et al., 2007). The homozygous mutation of *npas4l* significantly increased the number of *ins*:H2BGFP-positive cells during the β-cell regeneration period (Figures 1A-C). In addition, we observed a distinctive ectopic β-cell population in the mesenchymal region outside the pancreas in the *npas4l*^−/−^ group, an ectopic location that was very rarely observed in the sibling controls (including both wildtype siblings and heterozygous mutants). This ectopic population of β-cells contributed to the major increase in the number of *ins*:H2BGFP-positive cells during β-cell regeneration (Figure 1C). Moreover, the comparable and sparse numbers of *ins*:DsRed-positive cells in the controls and mutants indicate that the *npas4l* mutation did not enhance the survival of β-cells during the ablation (Figure 1A and B) because the extended maturation time of DsRed (Baird et al., 2000) restricted the detection of DsRed to the surviving β-cells.

**Figure 1.**
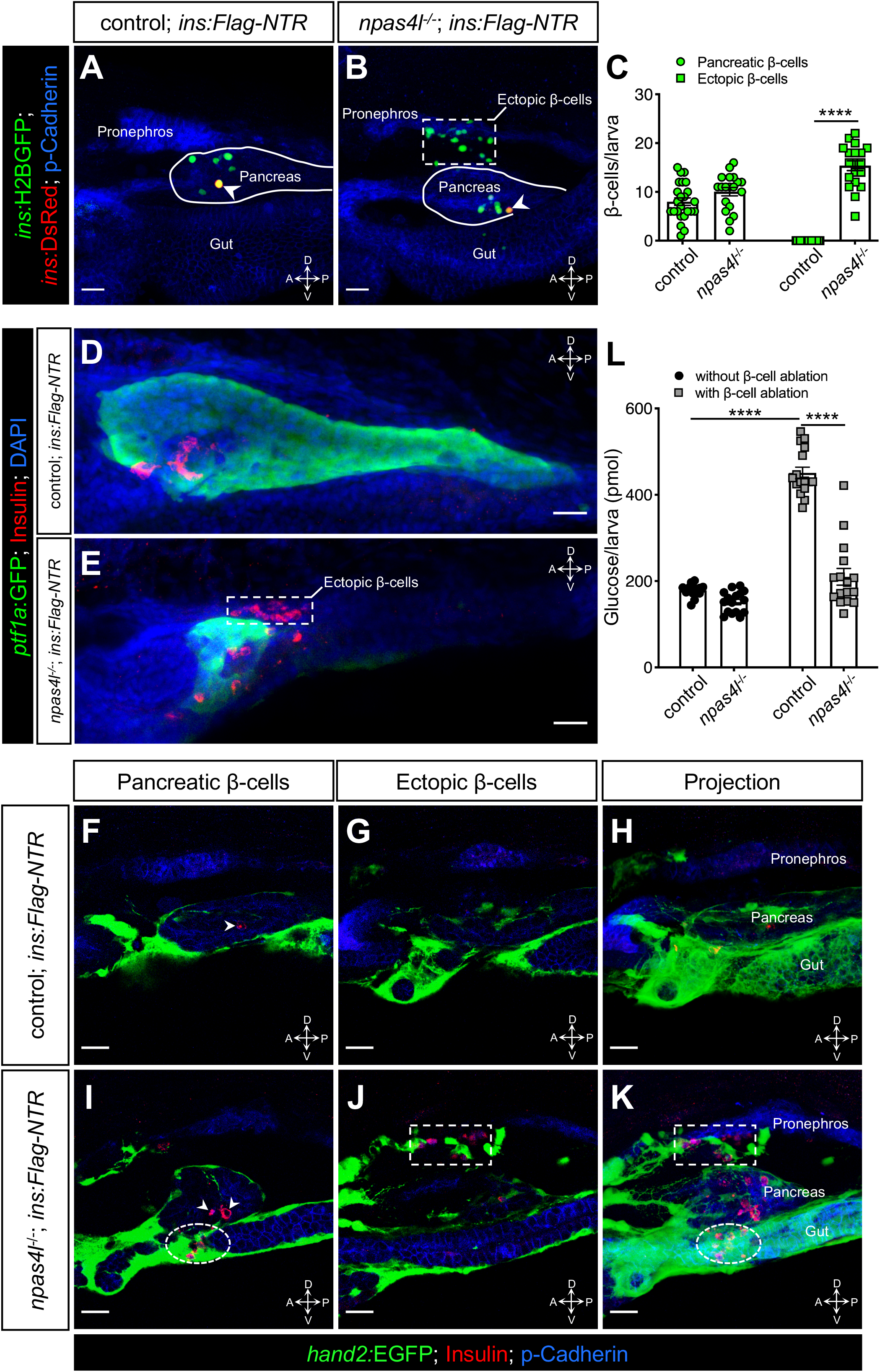
Ectopic β-cell formation and improved glucose control in *npas4l* mutants. (**A** and **B**) Representative confocal projections of the pancreas and the neighbouring tissues of control siblings and *npas4l*^−/−^ *Tg(ins:Flag-NTR);Tg(ins:H2BGFP;ins:DsRed)* zebrafish larvae at 3 dpf after β-cell ablation by MTZ at 1-2 dpf, displaying regenerated β-cells in green and older β-cells that likely survived the ablation in yellow overlap (arrowheads). The ectopic β-cells are indicated by the white dashed rectangle. Pancreata are outlined by the solid white lines. (**C**) Quantification of the pancreatic or ectopic β-cells per larva at 3 dpf. \*\*\*\**P* < 0.0001 (Šidák’s multiple comparisons test); *n* = 24 (control) and 19 (*npas4l*^−/−^). (**D** and **E**) Representative image projections of the pancreas and the neighbouring tissues in control siblings and *npas4l*^−/−^ *Tg(ins:Flag-NTR);Tg(ptf1a:GFP)* larvae at 3 dpf after β-cell ablation by MTZ at 1-2 dpf, displaying insulin-expressing β-cells in red and exocrine pancreas in green. The dashed rectangle indicates ectopic β-cells in the mesenchyme (**E**). (**F**-**K**) Representative images and projections of the pancreas and the neighbouring mesenchyme of control siblings and *npas4l*^−/−^ *Tg(ins:Flag-NTR);Tg(hand2:EGFP)* zebrafish larvae at 3 dpf after β-cell ablation by MTZ at 1-2 dpf, displaying β-cells in red and *hand2:*EGFP^+^ mesenchyme in green. White arrowheads point to β-cells in the pancreas (**F** and **I**). Dashed rectangles indicate the ectopic β-cells intermingling with the mesenchyme between the pronephros and the pancreas, without co-expressing insulin and *hand2:*EGFP (**J** and **K**). Selected area in dashed ovals shows the ectopic β-cells intermingling with the mesenchyme ventral to the pancreas (**I** and **K**). (**L**) Glucose levels of control siblings and *npas4l*^−/−^ larvae at 3 dpf after β-cell ablation at 1-2 dpf. Free-glucose levels of larvae without ablating the β-cells are shown as the baseline reference. \*\*\*\**P* < 0.0001 (Šidák’s multiple comparisons test); *n* = 80 larvae (16 groups of 5 pooled larvae) per data column. Quantification data are represented as the mean ± SEM. Scale bars = 20 μm. Anatomical axes: D (dorsal), V (ventral), A (anterior) and P (posterior).

To visualise the location of the ectopic β-cells better, we labelled the pancreas with *ptf1a:*GFP and observed not only a drastic reduction in the pancreas size (Figures 1D, E and Figure 1-figure supplement 1) but also the regeneration of β-cells clearly outside the *ptf1a*-expressing exocrine pancreas in *npas4l* mutants (Figure 1E). By labelling the mesenchyme with *hand2:*EGFP (Figure 1F-K), we further revealed that the majority of ectopic β-cells formed in *npas4l* mutants intermingled with *hand2:*EGFP-positive mesenchymal cells between the pronephros and the pancreas (Figures 1J and K). In addition, we occasionally observed ectopic β-cells intermingled with *hand2:*EGFP-positive mesenchymal cells ventral to the pancreas (Figures 1I and K). Although the ectopic β-cells were located among the mesenchymal cells, they did not express *hand2:*EGFP.

Additionally, we examined the *sst2:*RFP-positive δ-cell population in the *npas4l* mutants and revealed a small but significant increase outside the pancreas after δ-cell ablation (Figure 1-figure supplement 2), suggesting that the effect of homozygous *npas4l* mutation on ectopic endocrine cell formation is not limited to β-cells, albeit likely with a preference.

We further assessed the functionality and maturity of the ectopic β-cell population. We measured glucose levels in the control and *npas4l*^−/−^ groups with or without β-cell ablation to examine whether the newly formed β-cells could restore glucose to a normal level. Without β-cell ablation, the mutation of *npas4l* did not alter the glucose level, indicating that the *npas4l* mutation does not influence glucose homeostasis in the basal state (Figure 1L). After β-cell ablation, we observed an increased level of glucose in the sibling controls, while the homozygous mutation of *npas4l* resulted in a glucose level comparable to that of the controls without β-cell ablation, suggesting that the ectopic β-cells induced by the *npas4l* mutation contribute to restoring a physiological glucose level.

### The ectopic β-cells co-expressed insulin and endocrine markers in *npas4l* mutants

Next, we examined multiple pancreatic endocrine and β-cell markers, including Isl1, *neurod1*, *pdx1*, *mnx1*, *pcsk1* and *ascl1b* (the functional homolog to *Neurog3* in mammals), to validate the β-cell identity of the ectopic insulin-producing cells. The majority of ectopic β-cells co-expressed insulin and these markers during β-cell regeneration (Figure 2). The high co-expression of *pcsk1* (Figures 2R-S and Figure 2-figure supplement 1), which encodes an enzyme necessary for insulin biosynthesis, indicates that most of the β-cells in the ectopic population are likely functional. Consistent with preceding findings in pancreatic β-cells, not all ectopic β-cells expressed *ascl1b:*GFP (Figures 2V-W and Figure 1-figure supplement 1), which suggests that *ascl1b* works as a transient endocrine cell fate regulator (Flasse et al., 2013). In contrast with Isl1, *mnx1, pcsk1* and *ascl1b*, we observed lower co-expression levels of *neurod1* and *pdx1* in ectopic β-cells compared with the pancreatic population in *npas4l* mutants (Figure 2-figure supplement 1). In addition to the reduction in pancreas size (Figure 1-figure supplement 1), the *pdx1*-expressing pancreatic duct was also reduced in the *npas4l* mutant (Figure 2-figure supplement 2), indicating that the pancreas and its duct did not expand to form the ectopic β-cells. These observations together suggest that the pancreatic and ectopic β-cells are similar, yet they are two distinct β-cell populations.

**Figure 2.**
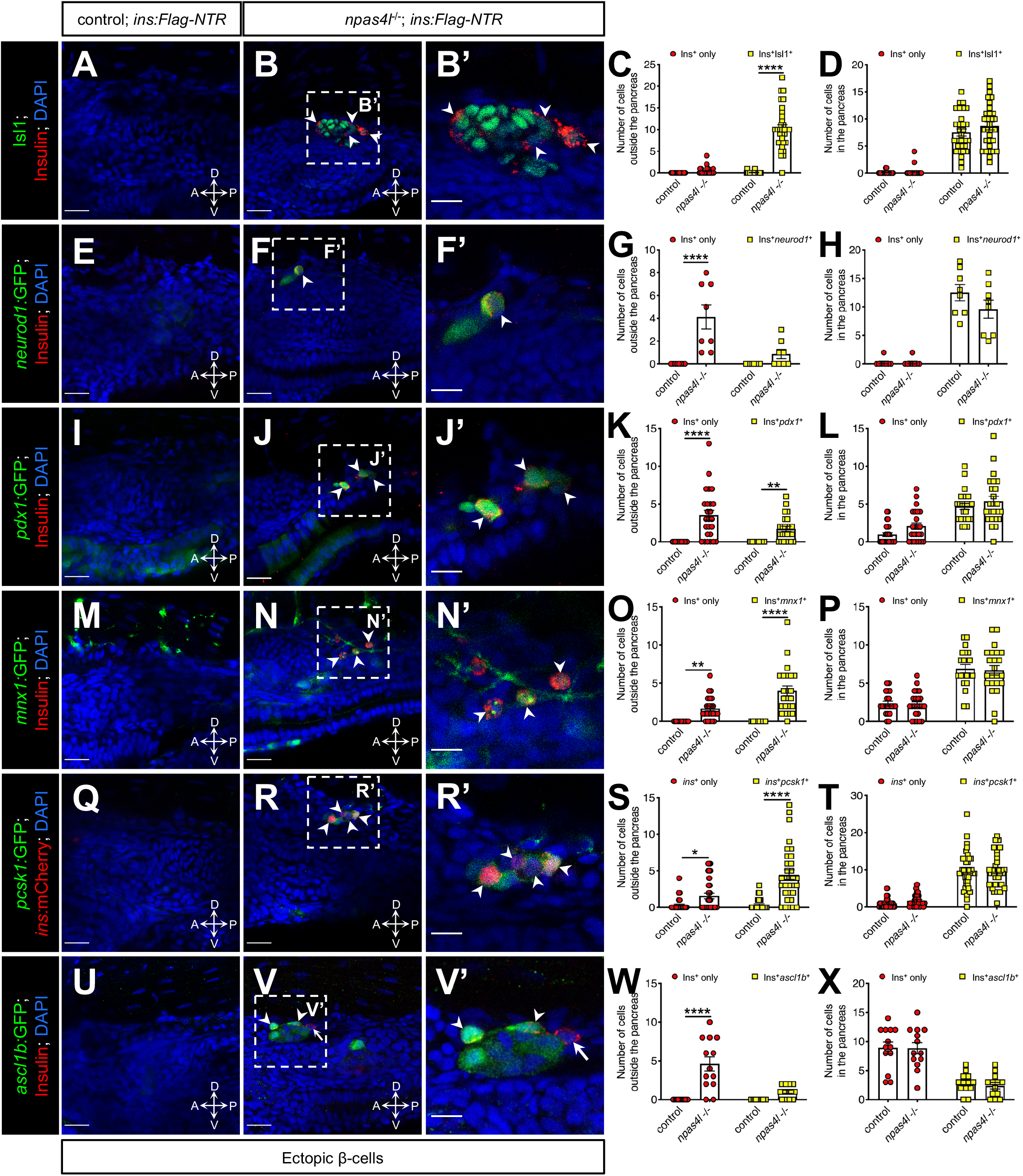
The ectopic β-cells co-expressed insulin and endocrine markers in *npas4l* mutants. Representative confocal images of the tissues adjacent to the pancreas of control siblings and *npas4l*^−/−^ *Tg(ins:Flag-NTR)* zebrafish larvae at 3 dpf after β-cell ablation by MTZ at 1-2 dpf, displaying cells expressing pancreatic endocrine cell markers Isl1 (**A**-**B’**), *neurod1* (**E**-**F’**), *pdx1* (**I**-**J’**), *mnx1* (**M**-**N’**), *pcsk1* (**Q**-**R’**) and *ascl1b* (**U**-**V’**) in green, and ectopic β-cells in red. Arrowheads point to ectopic β-cells that expressed corresponding markers. Arrows point to β-cells that did not express *ascl1b* (**V** and **V’**). **B’**, **F’**, **J’**, **N’**, **R’** and **V’** are magnified from the areas indicated by the white dashed square in **B**, **F**, **J**, **N**, **R** and **V** respectively. Quantification of β-cells with or without corresponding marker expression in the ectopic location (**C**, **G**, **K**, **O**, **S** and **W**) or in the pancreas (**D**, **H**, **L**, **P**, **T** and **X**) per larva at 3 dpf. \**P* = 0.0310, \*\**P* = 0.0039 and \*\*\*\**P* < 0.0001(Šidák’s multiple comparisons test); (**C** and **D**) *n* = 30 (control) and 31 (*npas4l*^−/−^); (**G** and **H**) *n* = 8 (control) and 8 (*npas4l*^−/−^); (**K** and **L**) *n* = 25 (control) and 24 (*npas4l*^−/−^); (**O** and **P**) *n* = 21 (control) and 23 (*npas4l*^−/−^); (**S** and **T**) *n* = 40 (control) and 32 (*npas4l*^−/−^); (**W** and **X**) *n* = 13 (control) and 13 (*npas4l*^−/−^). Data are represented as the mean ± SEM. Scale bars = 20 μm except **B’**, **F’**, **J’**, **N’**, **R’** and **V’** (10 μm). Anatomical axes: D (dorsal), V (ventral), A (anterior) and P (posterior).

### The ectopic β-cells in *npas4l* mutants and *etv2* morphants were of mesodermal origin

We have previously shown that *npas4l* expression is first initiated in the lateral plate mesoderm at the tailbud stage by *in situ* hybridization (Reischauer et al., 2016). In this study, we examined *npas4l* expression at 20 hpf, and found that *npas4l* was severely reduced in the lateral plate mesoderm in the *npas4l* mutants (Figure 3-figure supplement 1), whereas normal expression levels were observed in the tailbud and brain. The cells with reduced *npas4l* expression were still present in the lateral plate mesoderm as demonstrated by the embryos incubated overnight to further develop the *npas4l* expression signal (Figure 3-figure supplement 1B’). Because the ectopic β-cells induced by the *npas4l* mutation also resided in the mesenchymal region, and *npas4l* can act cell-autonomously to affect the hematopoietic and endothelial lineages (Parker & Stainier, 1999), we hypothesized that the ectopic β-cells originated from a mesodermal lineage.

**Figure 3.**
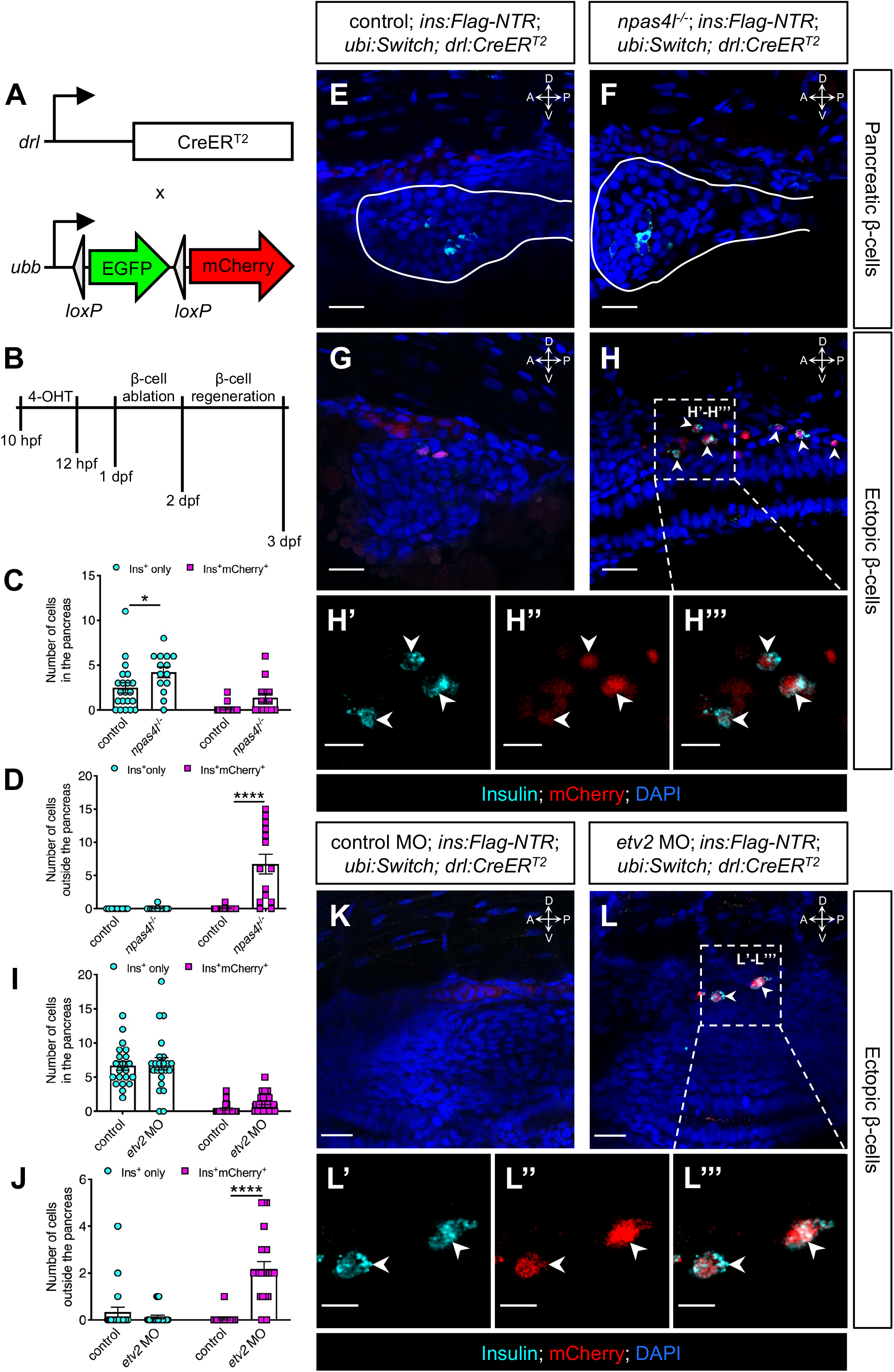
The ectopic β-cells in *npas4l* mutants and *etv2* morphants were of mesodermal origin. (**A**) Constructs of -*6.35drl:Cre-ER*^*T2*^ (*drl:CreER*^*T2*^) and -*3.5ubb:loxP-EGFP-loxP-mCherry* (*ubi:Switch*). Upon 4-OHT induction between 10-12 hpf, Cre-recombinase expressed by the *drl* promoter cleave the *loxP* sites to allow *ubb:mCherry* expression in the cells that once expressed *drl*. (**B**) Scheme for tracing the mesodermal lineage of ectopic β-cells in control siblings and *npas4l*^−/−^ *Tg(ins:Flag-NTR);Tg(ubi:Switch);Tg(drl:CreER^T2^)* zebrafish larvae. (**C**, **D**, **I** and **J**) Quantification of the pancreatic or ectopic β-cells with or without mesodermal lineage in *npas4l* mutants (**C** and **D**) or *etv2* morpholino (MO)-injected larvae (**I** and **J**) at 3 dpf. \**P* = 0.0227 and \*\*\*\**P* < 0.0001 (Šidák’s multiple comparisons test); (**C** and **D**) *n* = 21 (control) and 14 (*npas4l*^−/−^) or (**I** and **J**) *n* = 21 (control) and 23 (*etv2* MO). Data are represented as the mean ± SEM. (**E**-**H’’’** and **K**-**L’’’**) Representative confocal images of pancreatic (**E** and **F**) or ectopic β-cells (**G** and **H**) of control siblings and *npas4l*^−/−^, or ectopic β-cells in control or *etv2* MO-injected (**K** and **L**) *Tg(ins:Flag-NTR);Tg(ubi:Switch);Tg(drl:CreER^T2^)* zebrafish larvae at 3 dpf after β-cell ablation by MTZ at 1-2 dpf, displaying β-cells in cyan and lineage-traced cells derived from *drl*-expressing mesodermal cells in red. Pancreata are outlined by the solid white lines (**E** and **F**). Arrowheads point to ectopic β-cells derived from the mesoderm (**H**-**H’’’** and **L**-**L’’’**). Selected areas in dashed squares in **H** and **L** are magnified in split (**H’**, **H’’**, **L’** and **L’’**) and merged (**H’’’** and **L’’’**) channels, respectively. Scale bars = 20 μm (**E**-**H**, **K** and **L**) or 10 μm (**H’**-**H’’’** and **L’**-**L’’**). Anatomical axes: D (dorsal), V (ventral), A (anterior) and P (posterior).

To determine whether the mesoderm was the origin of the ectopic β-cells, we genetically traced the mesodermal cells using *drl:CreER*^*T2*^, a tamoxifen-inducible Cre transgene driven by a *drl* promoter (Mosimann et al., 2015). The spatial expression pattern of *drl* in the *npas4l* mutants resembled that in the sibling controls (Figure 3-figure supplement 2), suggesting that *npas4l* mutation did not induce any ectopic expression of *drl* to disrupt the lineage-tracing approach. Together with *ubb:loxP-EGFP-STOP-loxP-mCherry* (*ubi:Switch*) (Mosimann et al., 2011), the *drl*-expressing mesodermal cells would be labelled in red in *Tg(drl:CreER^T2^);Tg(ubi:Switch);Tg(ins:Flag-NTR)* (*drl*-tracing) zebrafish larvae after 4-hydroxytamoxifen (4-OHT) induction (Figure 3A). We treated the transgenic embryos with 4-OHT at 10-12 hours postfertilization (hpf). We chose to label the mesodermal cells during this period as neither endothelial/hematopoietic cells nor β-cells have developed at that stage, i.e. to exclude confounding effects of endothelial/hematopoietic cells or possible ectopic expression of the lineage tracer in the β-cells of the *npas4l* mutant. To ablate the β-cells, we incubated the 4-OHT-treated transgenic embryos in MTZ at 1-2 days postfertilization (dpf). We allowed the β-cells to regenerate for 30 hours before we fixed the larvae at 3 dpf for immunostaining (Figure 3B).

Immunostaining against insulin displayed a normal set of β-cells in the pancreas of the *drl*-tracing larvae with or without *npas4l* mutation after 30 hours of regeneration (Figures 3C, E and F). In line with the findings shown in Figure 1, the *npas4l* mutation induced the formation of ectopic β-cells in the mesenchymal region (Figures 3D, G and H). Furthermore, 98.9% of the ectopic β-cells in the mesenchymal region were mCherry-positive (Figures 3H’-H’’’), indicating that they derived from the *drl*-expressing mesodermal cells.

With a similar setting, we injected the *drl*-tracing embryos (without any *npas4l* mutation) with control or *etv2* morpholino at one-cell stage. Npas4l is essential for the expression of *etv2*, which is a key regulator of endothelial cell specification and vasculogenesis (Reischauer et al., 2016; Sumanas & Lin, 2006). Similar to *npas4l* mutation, knocking down *etv2* led to the formation of ectopic β-cells (Figures 3I-L’’’). The majority of the ectopic β-cells (94.3%) in *etv2* morphants was also lineage-traced back to the *drl*-expressing mesodermal cells, suggesting that the ectopic β-cell formation was also of mesodermal origin following *etv2* knockdown.

### The ectopic β-cells in *etv2* morphants derived from the *etv2*-expressing mesodermal lineage

To confirm the origin of the ectopic β-cell using a different lineage-tracing approach we generated *Tg(etv2:iCre)* zebrafish, which we then crossed into *Tg(ubi:Switch);Tg(ins:Flag-NTR)*, labelling *etv2*-expressing mesodermal and endothelial cells in red (Figure 4A). At the one-cell stage, we injected the *etv2*-tracing embryos with control or *etv2* morpholinos. After β-cell ablation by MTZ treatment at 1-2 dpf and β-cell regeneration for 30 hours ectopic β-cells formed in the *etv2* morphants, and 73.9% of the ectopic β-cells were labelled in red (Figures 4B-E’’’), illustrating that the *etv2*-expressing lineage gave rise to a significant portion of the ectopic β-cells.

**Figure 4.**
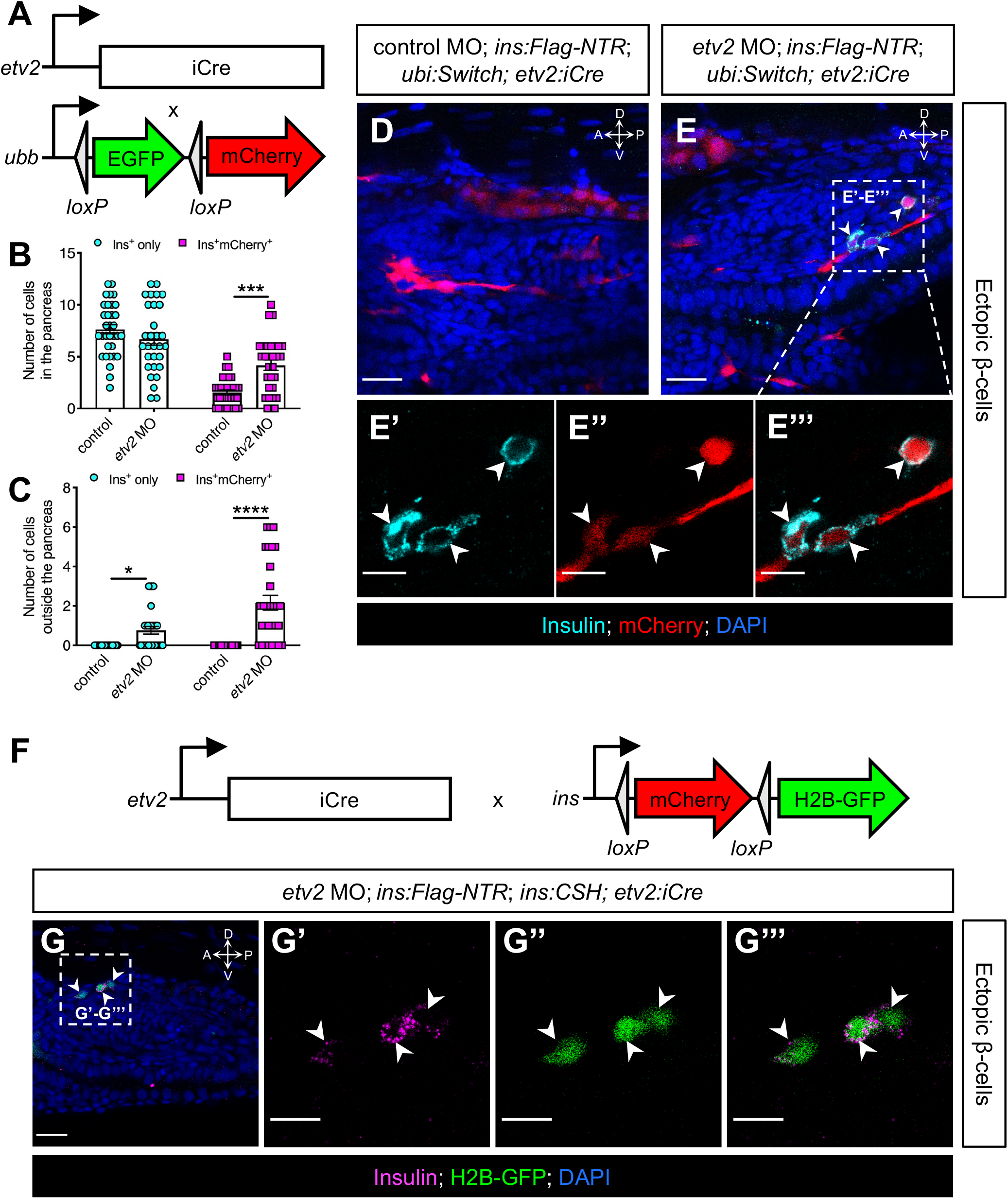
The ectopic β-cells in *etv2* morphants derived from the *etv2*-expressing mesodermal lineage. (**A**) Constructs of -*2.3etv2:iCre* (*etv2:iCre*) and -*3.5ubb:loxP-EGFP-loxP-mCherry* (*ubi:Switch*). (**B** and **C**) Quantification of the pancreatic or ectopic β-cells with or without *etv2*-positive mesodermal origin in control *or etv2* morpholino (MO)-injected larvae at 3 dpf. \**P* = 0.0169, \*\*\**P* = 0.0002 and \*\*\*\**P* < 0.0001 (Šidák’s multiple comparisons test); *n* = 33 (control) and 30 (*etv2* MO). Data are represented as the mean ± SEM. (**D**-**E’’’**) Representative confocal images of ectopic β-cells and *etv2*-positive lineage-traced cells in control or *etv2* MO-injected *Tg(ins:Flag-NTR);Tg(ubi:Switch);Tg(etv2:iCre)* zebrafish larvae at 3 dpf after β-cell ablation by MTZ at 1-2 dpf, displaying β-cells in cyan and lineage-traced cells derived from *etv2*-expressing mesodermal cells in red. The selected area in a dashed square in **E** is magnified in split (**E’** and **E’’**) and merged (**E’’’**) channels, respectively. (**F**) Constructs of -*2.3etv2:iCre* (*etv2:iCre*) and *ins:loxP-mCherry-loxP-H2B-GFP* (*ins:CSH*). (**G**-**G’’’**) Representative confocal images of ectopic β-cells derived from the *etv2*-expressing lineage in *etv2* MO-injected *Tg(ins:Flag-NTR);Tg(ins:CSH);Tg(etv2:iCre)* zebrafish larvae at 3 dpf after β-cell ablation by MTZ at 1-2 dpf, displaying β-cells in magenta and lineage-traced cells derived from *etv2*-expressing mesodermal cells in nuclear green. The selected area in a dashed square in **G** is magnified in split (**G’** and **G’’**) and merged (**G’’’**) channels, respectively. Scale bars = 20 μm (**D**, **E** and **G**) or 10 μm (**E’**-**E’’’** and **G’**-**G’’’**). Anatomical axes: D (dorsal), V (ventral), A (anterior) and P (posterior).

Moreover, we replaced *ubi:Switch* with *ins:loxP-mCherry-STOP-loxP-H2B-GFP* (*ins:CSH*) in the *etv2*-tracing zebrafish larvae to directly trace *insulin*-expressing cells originating from the *etv2*-expressing mesodermal lineage (Figure 4F). The co-localisation of insulin staining and the nuclear green tracer further confirms the mesodermal lineage of the ectopic β-cells (Figures 4G-G’’’).

Together, we used several different lineage-tracing models as well as two different loss of function models, i.e. using either the promoter of *drl* or *etv2* to drive Cre in either *naps4l* mutants or *etv2* morphants. This suggests that the ectopic β-cell formation is not restricted to the loss of a specific gene, but rather due to the absence of endothelial specification.

## Discussion

In this study, we first examined the role of blood vessels in β-cell regeneration in the *cloche* zebrafish mutant, which carries a homozygous *npas4l* mutation (Reischauer et al., 2016). We then unexpectedly revealed β-cells regenerating ectopically in the mesenchymal area. The ectopic β-cells were likely functional because they expressed several endocrine and β-cell markers including Isl1, *mnx1* and *pcsk1*, and were capable of restoring glucose levels during β-cell regeneration, although we do not know if they possess all the features of *bona fide* β-cells. Via *in situ* hybridization, lineage tracing and confocal microscopy, we successfully traced the origin of the ectopic β-cells to the mesodermal lineage. A recent study has reported the conversion of Etv2-deficient vascular progenitors into skeletal muscle cells, and highlighted the plasticity of mesodermal cell fate determination within the same germ layer (Chestnut et al., 2020). Our data demonstrated the plasticity of β-cell differentiation across the committed germ layers *in vivo*, i.e., switching from a mesodermal to an endodermal fate in a regenerative setting, while gastrulation and cell fate commitment in the germ layers are considered to be irreversible in development. Ectopic pancreata have been observed before, e.g. in *Hes1* mutant mice (Fukuda et al., 2006; Sumazaki et al., 2004), although that has been shown to be through an expansion of the pancreas rather than through changes in cell fate determination across organs or germ layers (Jorgensen et al., 2018). Our discovery is, to our knowledge, the first demonstration of ectopic β-cells with a mesodermal origin *in vivo*.

The mutated gene in the *cloche* mutant was named *npas4l* because its protein shares some homology with human NPAS4 (Reischauer et al., 2016). Although injecting either human *NPAS4* mRNA or zebrafish *npas4l* mRNA into zebrafish *cloche* mutant embryos at the one-cell stage could rescue the mutants, *Npas4* knockout mice are unlikely to share the same severe vascular and hematopoietic defects as zebrafish *npas4l* mutants because *Npas4* knockout mice survive to adulthood (Lin et al., 2008). This discrepancy suggests that other members of the mammalian NPAS protein family or other proteins may be functionally redundant with NPAS4 in vascular and hematopoietic development. Mammalian NPAS4 has been shown to have important cell-autonomous functions in β-cells (Sabatini et al., 2018; Speckmann et al., 2016). In zebrafish, *npas4a* is the main *npas4* paralog expressed in β-cells (Tarifeno-Saldivia et al., 2017), meaning that it is unlikely the phenotype we identified early in development in *npas4l* mutants is related to the functions of Npas4 in β-cells. Further studies on NPAS4, related bHLH transcription factors and ETV2 in mammals would elucidate whether inactivating such factors promotes β-cell formation with or without significantly perturbing the development of blood cells and vessels. We have shown that the enhanced differentiation potential in *npas4l* mutants is not limited to β-cell regeneration, which indicates that Npas4l may act as a gate for endodermal pancreatic cell fates in the mesoderm. Opening this gate in mesodermal cells may convert them to endodermal cells.

In summary, we have shown that the *npas4l* mutation or *etv2* knockdown induces ectopic regeneration of functional β-cells from the mesoderm. Our findings suggest a plasticity-potential of the mesodermal cells to differentiate into β-cells and other endodermal pancreatic cells (Figure 4-figure supplement 1). Further studies on the restriction of this plasticity would not only increase our understanding of the gating role of Npas4l and Etv2 in cell fate determination but also help to exploit an alternative source for β-cell regeneration.

## Methods

### Zebrafish

The following previously published mutant or transgenic zebrafish (*Danio rerio*) lines were used: *cloche*^*S5*^ (Field et al., 2003) as the *npas4l* mutant, *Tg(ins:Hsa.HIST1H2BJ-GFP;ins:DsRed)*^*s960*^ (Tsuji et al., 2014) abbreviated as *Tg(ins:H2BGFP;ins:DsRed)*, *Tg(ins:Flag-NTR)*^*s950*^ (Andersson et al., 2012), *Tg(ptf1a:GFP)*^*jh1*^ (Godinho et al., 2005), *Tg(hand2:EGFP)*^*pd24*^ (Kikuchi et al., 2011), *TgBAC(neurod1:EGFP)*^*nl1*^ (Obholzer et al., 2008), *Tg(−6.5pdx1:GFP)*^*zf6*^ (Huang et al., 2001), *Tg(mnx1:GFP)*^*ml2*^ (Flanagan-Steet et al., 2005), *Tg(pcsk1:eGFP)*^*KI106*^ (Lu et al., 2016), *TgBAC(ascl1b:EGFP-2A-Cre-ERT2)*^*ulg006Tg*^ (Ghaye et al., 2015) abbreviated as *Tg(ascl1b:GFP)*, *Tg(−3.5ubb:loxP-EGFP-loxP-mCherry)*^*cz1701*^ (Mosimann et al., 2011) abbreviated as *Tg(ubi:Switch)*, *Tg(−6.35drl:Cre-ERT2,cryaa:Venus)*^*cz3333*^ (Mosimann et al., 2015) abbreviated as *Tg(drl:CreER^T2^)* (a generous gift from Christian Mosimann), *Tg(sst2:NTR,cryaa:Cerulean)*^*KI102*^ (Lu et al., 2016) abbreviated as *Tg(sst2:NTR)*, *Tg(sst2:RFP)*^*gz19*^ (Li et al., 2009) and *Tg(insulin:loxP-mCherry-STOP-loxP-H2B-GFP; cryaa:Cerulean)*^*s934*^ (Hesselson et al., 2011), which is referred to *Tg(ins:mCherry)* in Figure 2 and *Tg(ins:CSH)* in Figure 4.

The *Tg(etv2:iCre;cryaa:Venus)*^*KI114*^ line was generated by the Tol2 transposon system and the construct was made by MultiSite Gateway Cloning (Invitrogen). The amplicon of the-*2.3etv2* promoter was synthesised from zebrafish genomic DNA with a forward primer 5’- TATAGGGCGAATTGggtaccTTCAGTAAGCAGACTCCTTCAATCA -3’ and a reverse primer 5’- AGCTGGAGCTCCAccgcggTTCGGCATACTGCTGTTGGAC -3’ by Phusion High-Fidelity DNA Polymerase (Thermo Scientific) as an insert for In-Fusion Cloning (Takara Bio) with p5E-MCS using restriction sites KpnI and SacII to yield p5E-*etv2*. Subsequently p5E-etv2, pME-iCre and p3E-polyA were used for the LR reaction with the destination vector to generate the construct *etv2:iCre*.

Males and females ranging in age from 3 months to 2 years were used for breeding to obtain new offspring for experiments. Individuals were sorted into the control sibling group (*npas4l*^+/+^ or *npas4l*^+/−^) and the homozygous *npas4l* mutant group (*npas4l*^−/−^) based on the characteristic pericardial oedema and blood-cell deficit. Zebrafish larvae were allocated into different experimental groups based on their phenotypes and genotypes in experiments involving *cloche* mutants. In morpholino knockdown experiments, zebrafish embryos were randomly assigned to each experimental condition for injection. Experimental procedures were performed on the zebrafish from 10 hpf to 3 dpf before the completion of sex determination and gonad differentiation. All zebrafish, except homozygous *npas4l* mutants and *etv2* morphants, appeared healthy and survived to adulthood. The homozygous *npas4l* mutants exhibited pericardial oedema, bell-shaped hearts and blood deficits, as previously reported (Stainier et al., 1995).The *etv2* morphants had similar phenotypes. All studies involving zebrafish were performed in accordance with local guidelines and regulations, and approved by regional authorities.

### Chemical ablation of β- and δ-cells

As in our previous report (Schulz et al., 2016), we ablated β-cells and δ-cells by incubating the β-cell ablation model *Tg(ins:Flag-NTR)* zebrafish and the δ-cell ablation model *Tg(sst2:NTR)* zebrafish in E3 medium supplemented with 10 mM metronidazole (MTZ, Sigma-Aldrich), 1% DMSO (VWR) and 0.2 mM 1-phenyl-2-thiourea (Acros Organics) for 24 h from 1 to 2 dpf.

### Microinjection of morpholinos

Standard control morpholino (5’-CCTCTTACCTCAGTTACAATTTATA-3’) and *etv2* morpholino (5’-CACTGAGTCCTTATTTCACTATATC-3’) (Sumanas & Lin, 2006) were purchased from Gene Tools, LLC and 4ng of each was injected into one-cell stage zebrafish embryos.

### Lineage tracing by tamoxifen-inducible Cre recombinase

To genetically trace the mesodermal lineage, we treated *Tg(ins:Flag-NTR);Tg(ubi:Switch);Tg(drl:CreER^T2^)* zebrafish embryos with 10 μM 4-OHT (Sigma-Aldrich) in E3 medium in 90-mm Petri dishes, with approximately 60 individuals per dish, from 10 to 12 hpf. Upon induction by 4-OHT, cytoplasmic CreER^T2^ would be translocated to the nucleus to excise the loxP-flanked EGFP to enable mCherry expression in *drl*-expressing cells and their descendants, indicating a mesodermal lineage.

### Sample fixation for immunostaining

Before fixing the zebrafish larvae, we confirmed the presence of the transgenes by determining the corresponding fluorescent markers and subsequently examined them under a widefield fluorescence microscope LEICA M165 FC (Leica Microsystems). We then euthanized the zebrafish larvae with 250 mg/L tricaine (Sigma-Aldrich) in E3 medium followed by washing in distilled water three times. We fixed the samples in 4% formaldehyde (Sigma-Aldrich) in PBS (ThermoFisher Scientific) at 4 °C overnight. After washing away the fixative with PBS three times, we removed the skin and crystallized yolk of the zebrafish larvae by forceps under the microscope to expose the pancreas and mesenchyme for immunostaining.

### Immunostaining and confocal imaging

As in our previous report (Liu et al., 2018), we started immunostaining by incubating the zebrafish samples in blocking solution (0.3% Triton X-100, 4% BSA and 0.02% sodium azide from Sigma-Aldrich in PBS) at room temperature for one hour. We then incubated the samples in blocking solution with primary antibodies at 4 °C overnight. After removing the primary antibodies, we washed the samples with washing buffer (0.3% Triton X-100 in PBS) eight times at room temperature for two hours. Afterwards, we incubated the samples in blocking solution with fluorescent dye-conjugated secondary antibodies and the nuclear counterstain DAPI (ThermoFisher Scientific) if applicable at 4 °C overnight. Next, we removed the secondary antibodies and nuclear counterstain and washed the samples with washing buffer eight times at room temperature for two hours. The following primary antibodies were used: anti-GFP (1:500, Aves Labs, GFP-1020), anti-RFP (1:500, Abcam, ab62341), anti-insulin (1:100, Cambridge Research Biochemicals, customised), anti-pan-cadherin (1:5000, Sigma, C3678) and anti-islet-1-homeobox (1:10, DSHB, 40.3A4 supernatant).

Before confocal imaging, we mounted the stained samples in VECTASHIELD Antifade Mounting Medium (Vector Laboratories) on microscope slides with the pancreas facing the cover slips. We imaged the pancreas and the neighbouring mesenchyme of every zebrafish sample that we had mounted with the confocal laser scanning microscopy platform Leica TCS SP8 (Leica Microsystems).

### Determination of free glucose level in zebrafish

To collect the samples, we washed the zebrafish larvae in PBS and transferred them to individual tubes, with 5 larvae per tube, for snap freezing in liquid nitrogen. Afterwards, we added a 5-mm stainless steel bead (QIAGEN) and 100 μl of PBS to each tube and lysed the samples by TissueLyser II (QIAGEN) at 4 °C for 2 min. After centrifugation, we transferred the supernatant to another tube for further analysis.

We employed the Glucose Colorimetric/Fluorometric Assay Kit (BioVision) to measure the free glucose level in the zebrafish larvae according to the manufacturer’s protocol. First, we prepared a glucose standard at 0, 0.1, 0.2, 0.4 and 0.8 nmol in 25 μl of glucose assay buffer in a 96-well microplate for the standard curve. We then transferred 5 μl of the sample supernatant from each sample tube together with 20 μl of glucose assay buffer to the microplate. Subsequently, we prepared the glucose reaction mix consisting of 24.8 μl of glucose assay buffer, 0.1 μl of glucose probe and 0.1 μl of glucose enzyme mix for each reaction. After adding 25 μl of glucose reaction mix to each reaction, we incubated the microplate at 37 °C in the dark for 30 min. Finally, we measured the fluorescence intensity emitted from the reactions with the FLUOstar OPTIMA microplate reader (BMG LABTECH) at Ex/Em = 535/590 nm.

### Whole-mount in situ hybridization

Zebrafish embryos at 10 and 20 dpf were fixed with 4% paraformaldehyde in PBS at 4 °C overnight. Whole-mount *in situ* hybridization was performed according to the method in a previous report (Thisse & Thisse, 2008). Probes against *npas4l* and *drl* were synthesised from PCR-products using bud-stage zebrafish cDNA, Phusion High-Fidelity DNA Polymerase (Thermo Scientific) and primer pairs 5’- ACTCGGGCATCAGGAGGATC-3’ plus 5’- (CCTAATACGACTCACTATAGGG)GACACCAGCATACGACACACAAC-3’ for *npas4l*, and 5’- ATGAAGAATACAACAAAACCC-3’ plus 5’- (CCTAATACGACTCACTATAGGG)TGAGAAGCTCTGGCCGC-3’ for *drl*, respectively, T7 was employed for transcription, and digoxigenin (Roche) was used for labelling. To genotype the *npas4l* mutants, PCR was performed using gDNA from the imaged samples and primers 5’- TTCCATCTTCTGAATCCTCCA-3’ plus 5’- GGACAGACCCAGATACTCGT-3’ at the conditions previously reported (Reischauer et al., 2016). The PCR products were then sent for sequencing with a primer 5’- TTTCTGCCGTGAATGGATGTG-3’ (Eurofins Genomics).

### Statistical analysis

Similar experiments were performed at least two times independently. The number of cells in the confocal microscopy images were all quantified manually with the aid of the Multipoint Tool from ImageJ. Data were then analysed with Prism (GraphPad). Statistical analyses were carried out by two-tailed *t*-tests when two groups were analysed and by ANOVA when more than two groups were analysed. We have presented the results as the mean values ± SEM and considered *P* values ≤ 0.05 to be statistically significant. The *n* number represents the number of zebrafish individuals in each group of each experiment.

## Acknowledgments

Research in the lab of O.A. was supported by funding from the European Research Council under the Horizon 2020 research and innovation programme (grant n° 772365); the Swedish Research Council; the Novo Nordisk Foundation; Ragnar Söderberg’s Foundation; and the Strategic Research Programmes in Diabetes, and Stem Cells & Regenerative Medicine at the Karolinska Institutet. This work was also supported by an EFSD/Lilly Fellowship awarded to K.C.L.

## Figure Supplements Legends

**Figure 1-figure supplement 1.**
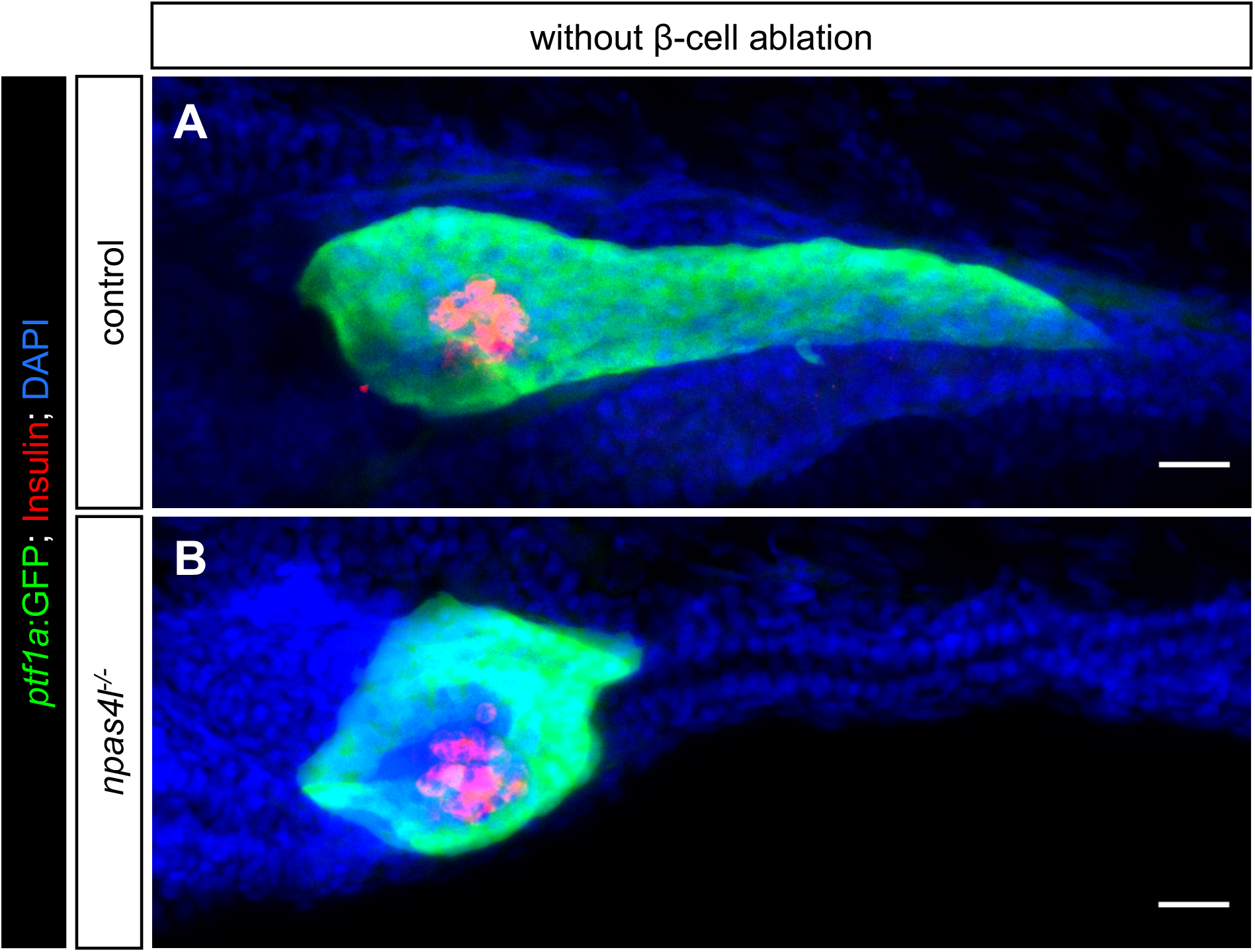
Mutation of *npas4l* suppressed the development of exocrine pancreas. (**A** and **B**) Representative image projections of the pancreas in control siblings and *npas4l*^−/−^ *Tg(ptf1a:GFP)* larvae, without carrying the *ins:Flag-NTR* transgene, i.e. during regular development. These larvae were controls for the β-cell ablation, i.e. still treated with MTZ from 1-2 dpf (not leading to β-cell ablation due to the absence of *ins:Flag-NTR*). Insulin-expressing β-cells are displayed in red and exocrine pancreas in green. Scale bars = 20 μm.

**Figure 1-figure supplement 2.**
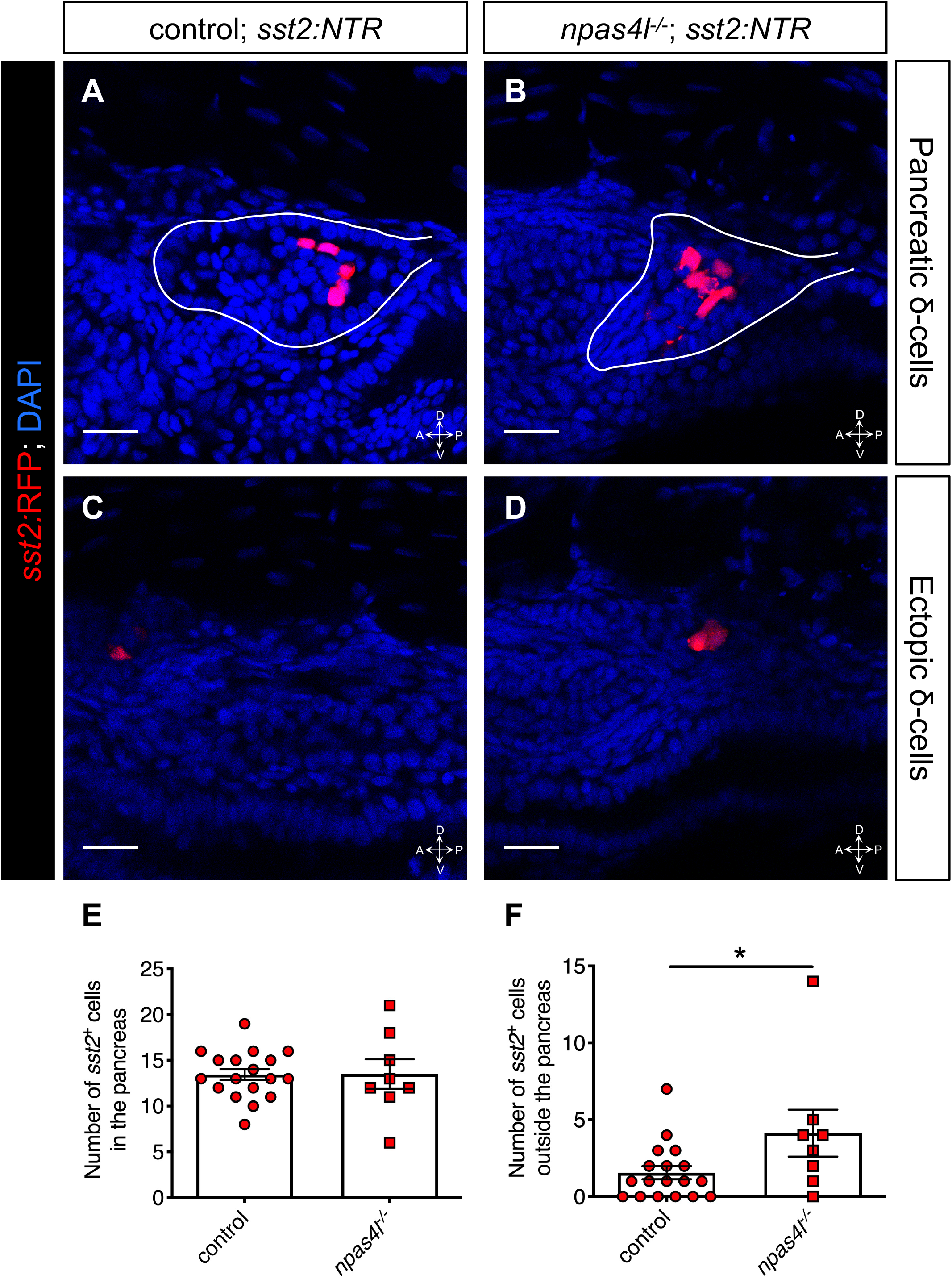
Mutation of *npas4l* mildly induced ectopic *sst2*-expressing δ-cells. (**A**-**D**) Representative images of pancreatic δ-cells (**A** and **B**) and ectopic δ-cells (**C** and **D**) of control siblings and *npas4l*^−/−^ *Tg(sst2:NTR);Tg(sst2:RFP)* zebrafish larvae at 3 dpf after δ-cell ablation by MTZ at 1-2 dpf; δ-cells are shown in red. Solid white lines outline the pancreata. (**E** and **F**) Quantification of the *sst2:*RFP^+^ δ-cells in the pancreas (**E**) or in the mesenchyme outside the pancreas (**F**) per larva at 3 dpf. * *P* = 0.0479 (Mann-Whitney test); *n* = 18 (control) and *n* = 8 (*npas4l*^−/−^). Data are represented as the mean ± SEM. Scale bars = 20 μm. Anatomical axes: D (dorsal), V (ventral), A (anterior) and P (posterior).

**Figure 2-figure supplement 1.**
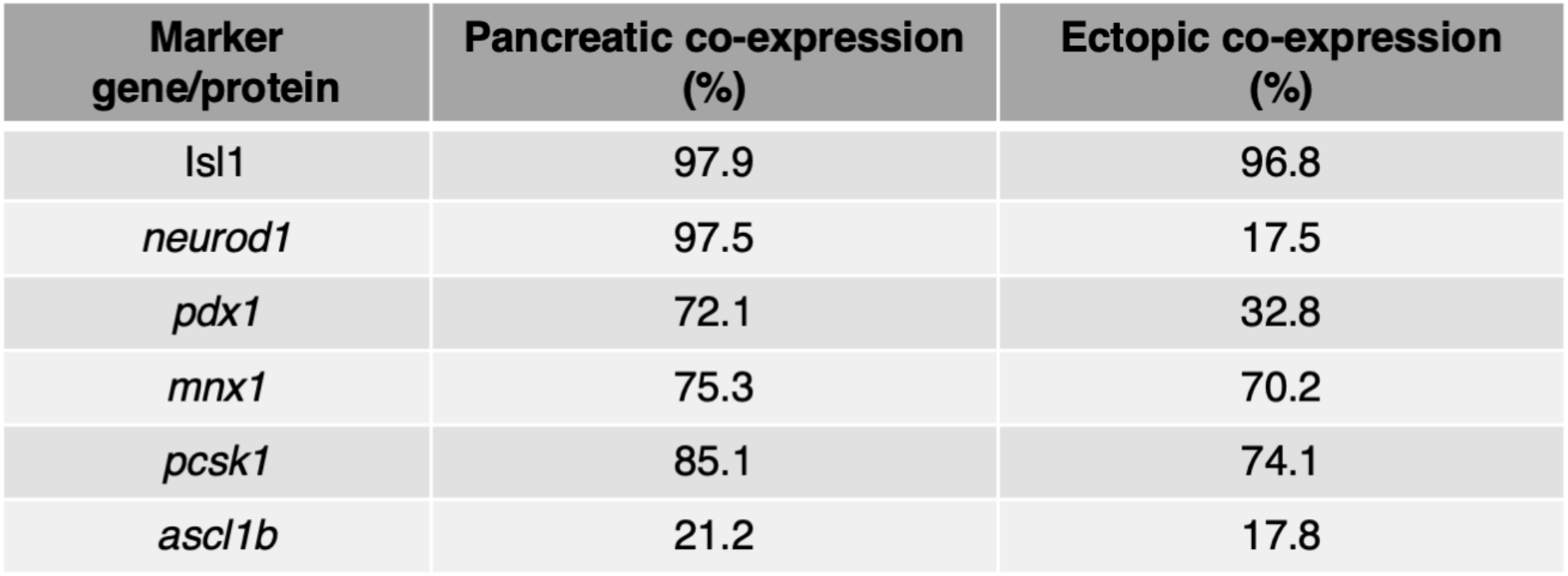
Percentages of pancreatic or ectopic cells co-expressing insulin and corresponding marker gene or protein in *npas4l* mutants.

**Figure 2-figure supplement 2.**
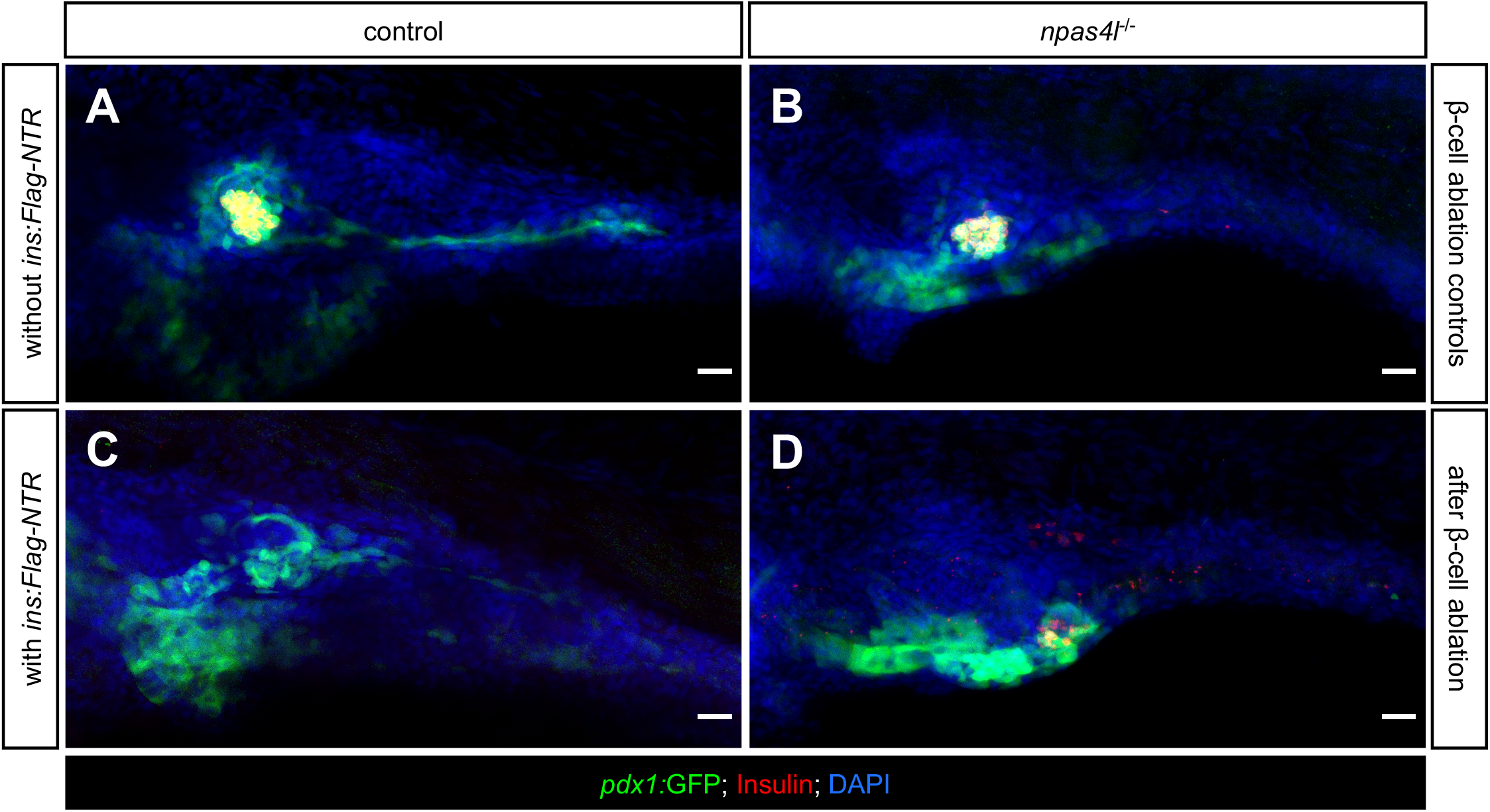
Mutation of *npas4l* inhibited *pdx1*-expressing pancreatic duct formation. (**A**-**D**) Representative image projections of the pancreas and the surrounding tissues in control siblings and *npas4l*^−/−^ *Tg(pdx1:GFP)* zebrafish larvae without (**A** and **B**) or with (**C** and **D**) ins*:Flag-NTR* expression at 3 dpf after treatment with MTZ at 1-2 dpf. Hence **A** and **B** were β-cell ablation controls while the β-cells in **C** and **D** had been ablated. Insulin-expressing β-cells are displayed in red and *pdx1*-expressing cells (including pancreatic ductal cells and upper intestine) in green. The dashed rectangle indicates that the ectopic β-cells were distant from the *pdx1*-expressing pancreatic ducts. Scale bars = 20 μm.

**Figure 3-figure supplement 1.**
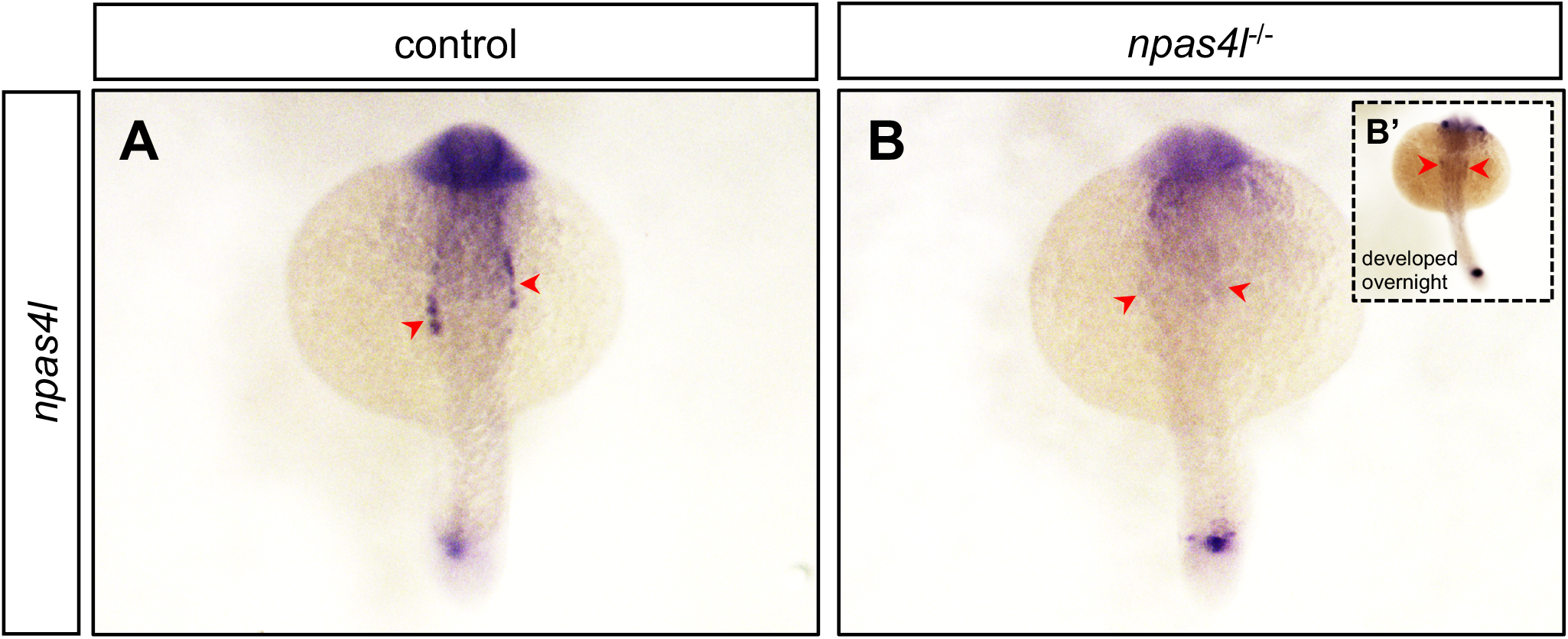
Cell population with reduced *npas4l* expression remains in the lateral plate mesoderm before β-cell ablation. (**A** and **B**) Representative images of *in situ* hybridization against *npas4l* expression in control siblings and *npas4l*^−/−^ zebrafish embryos at 20 hpf after a 45-minute incubation to develop the expression signal. Inset **B’** displays a representative *npas4l*^−/−^ zebrafish embryo incubated overnight to further develop the signal. Red arrowheads point to lateral plate mesoderm expressing *npas4l*.

**Figure 3-figure supplement 2.**
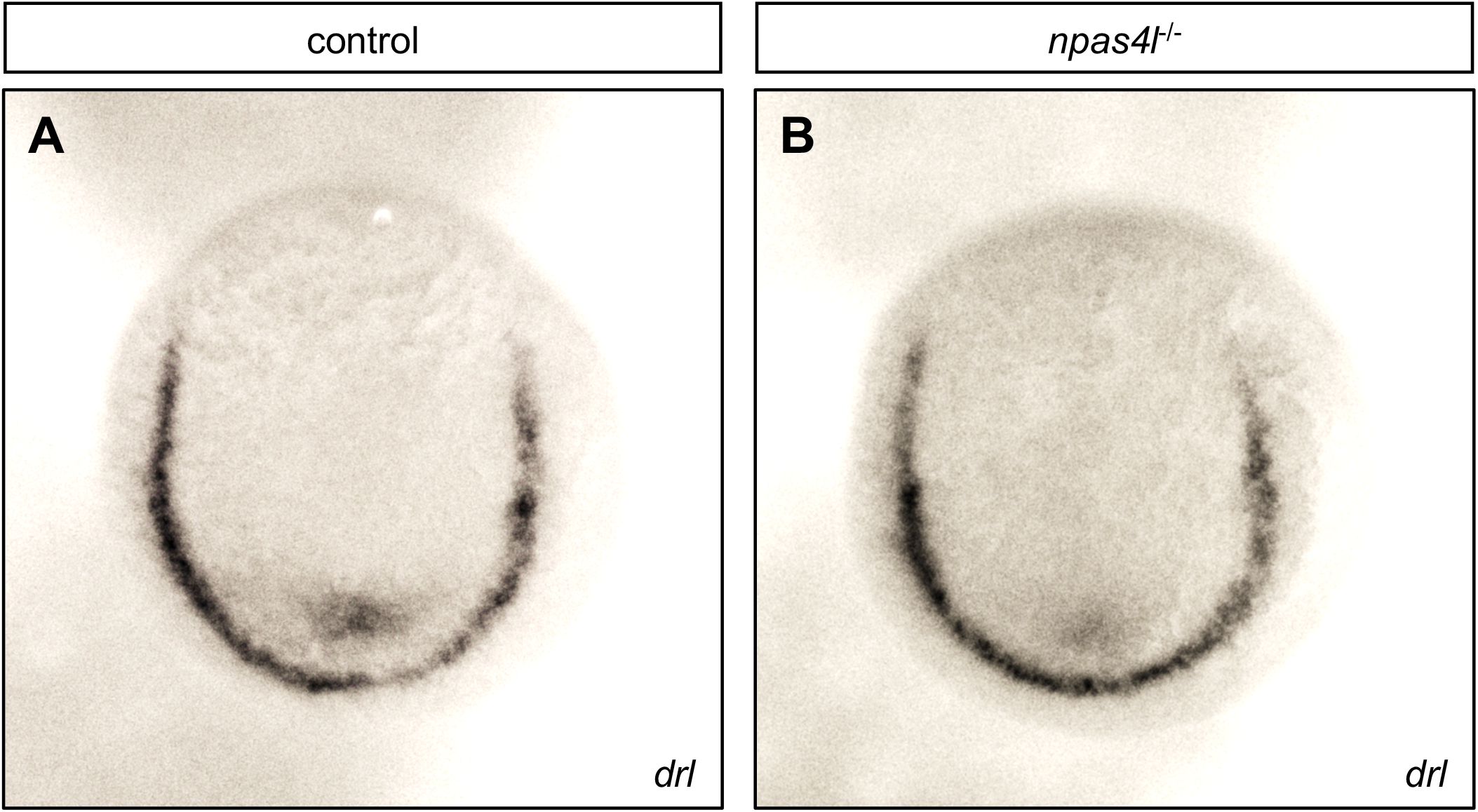
The *npas4l* mutant did not display altered expression of *drl* in the lateral plate mesoderm. (**A** and **B**) Representative images of *in situ* hybridization against *drl* expression in control siblings and *npas4l*^−/−^ zebrafish embryos at 10 hpf.

**Figure 4-figure supplement 1.**
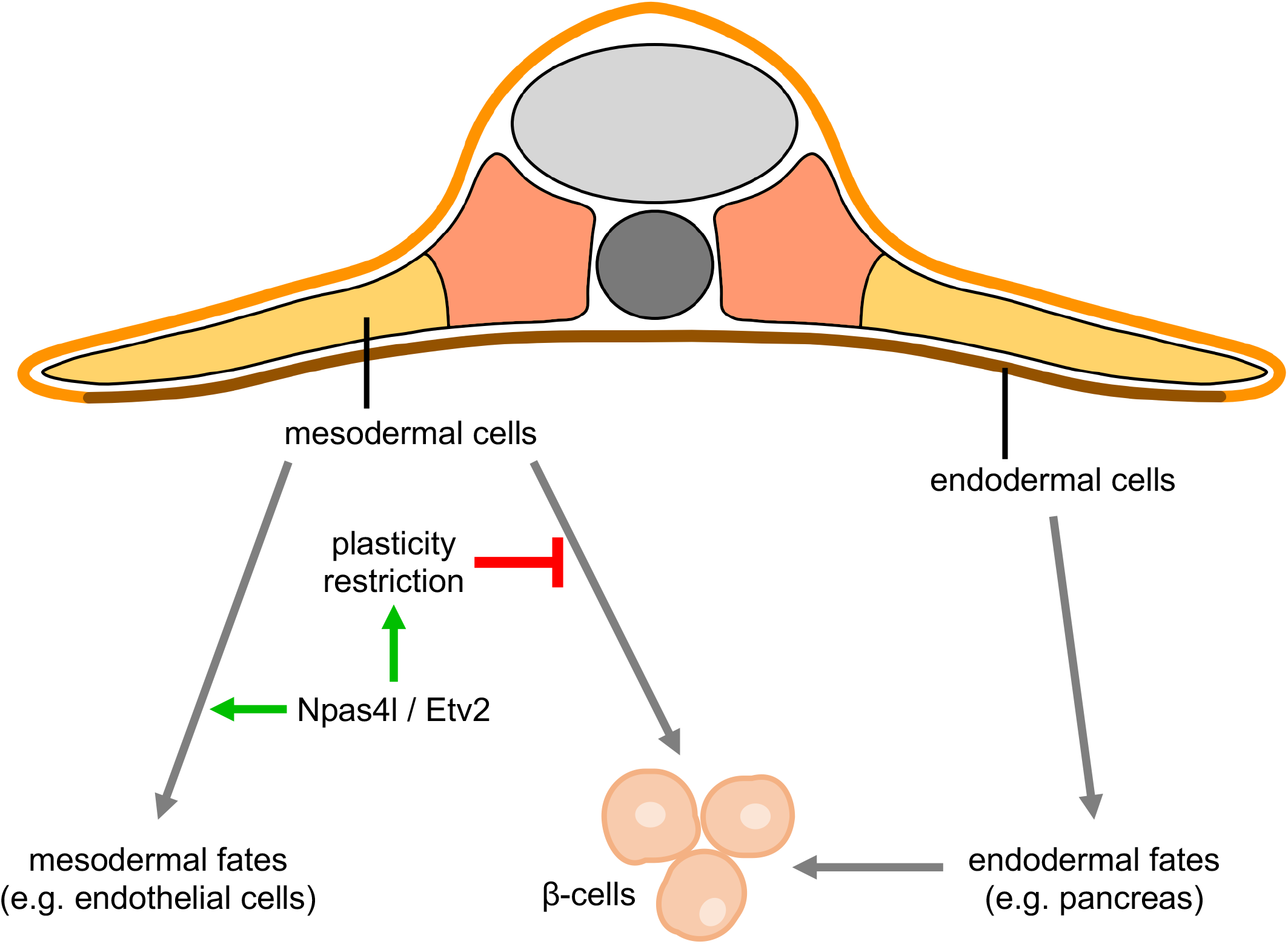
Npas4l/Etv2 restricts the plasticity of the mesoderm. Mesoderm and endoderm normally follow Waddington’s landscape model to further differentiate into cells with mesodermal fates and endodermal fates, respectively, during development. However, mutating *npas4l* or knocking down *etv2* not only abolishes the endothelial specification but also induces plasticity of mesodermal cells to enable their differentiation to β-cells, δ-cells and perhaps other endodermal pancreatic cells across the germ layer border.

